# Large Language Models for Pathway Curation: A Preliminary Investigation

**DOI:** 10.1101/2024.04.26.591413

**Authors:** Nikitha Karkera, Nikshita Karkera, Mahanash Kumar, Samik Ghosh, Sucheendra K. Palaniappan

## Abstract

The pathway curation task involves analyzing scientific literature to identify and represent cellular processes as pathways. This process, often time-consuming and labor-intensive, requires significant curation efforts amidst the rapidly growing biomedical literature. Natural Language Processing (NLP) offers a promising method to automatically extract these interactions from scientific texts. Despite immense progress, there remains room for improvement in these systems. The emergence of Large Language Models (LLMs) provides a promising solution for this challenge. Our study conducts a preliminary investigation into leveraging LLMs for the pathway curation task. This paper first presents a review of the current state-of-the-art algorithms for the pathway curation task. Our objective is to check the feasibility and formulate strategies of using these LLMs to improve the accuracy of pathway curation task. Our experiments demonstrate that our GPT-3.5 based fine-tuned models outperforms existing state-of-the-art methods. Specifically, our model achieved a 10 basis point improvement in overall recall and F1 score compared to the best existing algorithms. These findings highlight the potential of LLMs in pathway curation tasks, warranting further research and substantial efforts in this direction.

**Keypoints/Objectives:**

- Study evaluates the feasibility of using Large Language Models (LLMs) for pathway curation in scientific literature.
- Using GPT-3.5 based fine tuned models for pathway curation, we compare its performance with existing methods, focusing on precision, recall and F1 score metrics.
- Emphasize the promise and need for further research on using LLMs for pathway curation.

## Introduction

Pathway curation involves collecting and organizing data about biological pathways, crucial for understanding cellular processes and interactions. In our context, pathways represent a series of interconnected molecular events or interactions that collectively govern specific biological functions or processes (1). These functions could range from basic cellular activities to complex physiological responses. Curating these pathways requires integration of data from various sources, including experimental studies and scientific literature. Researchers often use these information to carefully assemble information about genes, proteins, and small molecules, and other relevant players along with their roles and relationships within the intricate web of cellular signalling and Regulation (2, 3). Significant efforts have been made in the detailed mapping of process or disease-specific pathways. Recent notable efforts include a comprehensive map of lipid mediator metabolic pathways(4), Alzheimer’s disease (5), Parkinson’s disease (6), and the influenza A virus replication cycle (7). Human curation, while accurate, is extremely time-consuming and cannot keep pace with the increase in the number of publications (8). There is a need to move towards automation of the extraction and interpretation which will significantly speed up the process and reduces the likelihood of human error or oversight. This not only make the process more efficient but also allows for the timely updating and expanding of pathway databases.

The goal of the Pathway Curation task is to using natural language processing (NLP) techniques to automatically extract and interpret information about these processes directly from scientific literature. The task that was defined in the BioNLP Shared Task 2013 (9), focuses on evaluating the applicability of event extraction systems to assist in the curation, evaluation, and maintenance of bio-molecular pathway models. Event extraction in our context refers to the process of identifying and characterizing biological events, such as gene expression, phosphorylation, complex formation from text. It further comprises of the following sub problems: 1) identifying relevant entities (like genes or proteins), 2) determining the relationships or interactions between these entities and 3) classifying these interactions into predefined categories. Over the years, many algorithms and platforms have been proposed to tackle the problem with varying degree of success (10)

The primary objective of this study is to investigate the effectiveness of using Large Language Models (LLMs) for the pathway curation task. Utilizing the sophisticated natural language understanding and generative capabilities of LLMs, we hope to surpass the current benchmarks in pathway curation efficiency and accuracy.

In terms of the organization of the paper, the next section describes major related work for tackling the pathway curation task, following this we will describe the data set which is used for fine tuning the large language models (LLMs). The following sections introduces our pipeline and results in detail.

### Literature Survey

There are multiple algorithms and systems that have been used for pathway curation. Notable among these is the EventMine system developed by the National Centre for Text Mining (NaCTeM)(11), which employs a one-vs-rest support vector machine (SVM) for event extraction. In terms of performance, accuracy ranges range approximately from 60% to 70% for non-nested events such as SIMPLE, around 40% for nested events like Regulation, and about 30% for modification events (e.g., MOD).

Another significant system is the Turku Event Extraction System (TEES) (9). Introduced in 2009, TEES leverages a unified graph representation and stepwise SVM-based machine learning for even extraction. TEES 1.0 achieved an F-score of 51.95%. Furthermore, TEES 2.2 (12) marked a significant advancement by automating the learning of annotation rules for various event types. This version stood out at BioNLP’13. The integration of the scikit-learn machine learning library into TEES 2.2 brought in a wider array of machine learning methods and enabled the analysis of feature importances in the TEES feature sets. Systems like EventMine (13), TEES SVM (14), DeepEventMine (15) were developed later on with DeepEventMine being the latest system with highest precision of 64.12% and TEES CNN with highest F-score of 58.31%.

Another approach leverages rule-based literature mining for extracting pathway information from text, particularly using curated pharmacokinetic (PK) and pharmacodynamic (PD) pathways in PharmGKB (16). This system achieved an F-measure of 63.11% for entity extraction and 34.99% for event extraction, providing a distinct point of comparison against systems like NaCTeM and TEES.

Several noteworthy efforts in the field have been made, including the work of (17), who implemented a sequence labeling strategy, and (18), who focused on attention mechanisms. (19) introduced a joint framework for document-level biomedical event extraction, employing a dependency-based Graph Convolutional Network (GCN) for local context and a hypergraph for global context. In their study, (20) provided a comprehensive evaluation of various datasets using the TEES 2.2 system. More recently, (21) employed multiple natural language processing systems alongside the Integrated Network and Dynamical Reasoning Assembler (INDRA), while (22) compared human and machine-curated HCM molecular mechanisms models using the INDRA system. Furthermore,

(23) explored a multi-turn question answering approach with BERT models, a method similarly adopted by (24). Each system and approach contributes uniquely to the evolving landscape of pathway curation.

The domain of Large Language Models (LLMs) have seen significant progress in the past year(25). These models use deep learning techniques to analyze, generate, and understand human language at scale having trained on vast datasets to accurately model linguistic patterns and context. For instance, GPT-3.5 (26), a state-of-the-art LLM model by OpenAI, has garnered acclaim for its exceptional performance for a variety of NLP tasks. LLMs ability to process and synthesize complex information makes them particularly suitable for tasks requiring high levels of language comprehension, such as the pathway curation task. They can be fine-tuned for specific tasks, tailoring their capabilities to specialized requirements. For the pathway curation task, we fine-tune the GPT-3.5 model using curated training data previously available for this task. More details of the training data set and the fine tuning method is discuss in further sections.

### Pathway curation dataset

#### Data Representation

For the pathway curation task, data representation is key and it involves categorizing and detailing various entities and events within biological pathways. The pathway data is represented using the ST format that was made popular in the BioNLP community through various shared tasks (9, 27, 28). The main components of the format which are relevant to our work is detailed below.

#### Entities

There are four key entity types in pathway curation: SIMPLE CHEMICAL, GENE OR GENE PRODUCT, COMPLEX, and CELLULAR COMPONENT. Each entity is annotated in the text with specific references to relevant biological databases. SIMPLE CHEMICAL refers to the ChEBI resource, GENE OR GENE PRODUCT to gene and protein databases, COMPLEX to complex-specific databases, and CELLULAR COMPONENT to the Gene Ontology’s cellular component. We refer the readers to (9) for more details.

#### Events

The event annotation marks references to biological reactions, processes, and associations. For the details of each event type, we refer the reader to (9) for more details. Each event argument, whether an entity or another event, is specified with a defined role. This setup facilitates detailed representation and analysis of complex biological interactions within pathway models. These roles include “Theme”, “Cause”, location-based roles such as “AtLoc”, “FromLoc”, and “ToLoc”, the “Site” of modifications, and the “Participant” in broader processes. Each of these role types plays a unique part in describing interactions in biological pathways. The following sections delve into each of these role types in detail, offering insights into their significance and application in pathway curation.

- Theme - entity/event that undergoes the effects of the event. For example, the entity that is transcribed in a TRANSCRIPTION event or transported in a TRANSPORT event.
- Cause - entity/event that is causally active in the event. Marks, for example, “P1” in “P1 inhibits P2 expression”.
- AtLoc,FromLoc,ToLoc - location in which the Theme entity of a LOCALIZATION event is localized (At) in LOCALIZATION events not involving movement or is transported (or moves) from/to (From/To) in LOCALIZATION and TRANSPORT events involving movement.
- Site - site on the Theme entity that is modified in the event. Can be specified for modification events such as PHOSPHORYLATION.
- Participant - general role type identifying an entity that participates in some underspecified way in a high-level process. Only applied for the PATHWAY type.

### Data Insights

The data used for this paper is from the Pathway Curation data of the various relevant BioNLP-ST (9) tasks.

#### A. Structure of data

As mentioned before, for this paper, we used the pathway curation dataset from BioNLP-ST 2013. For every abstract (scientific text), the raw data is divided into three files as shown in Fig. 2 : 1st file (.txt) contains the text abstract of scientific publication, 2nd file (.a1) contains all the entities in the abstract and the 3rd file (.a2) contains the events, along with their roles.

**Fig. 1.**
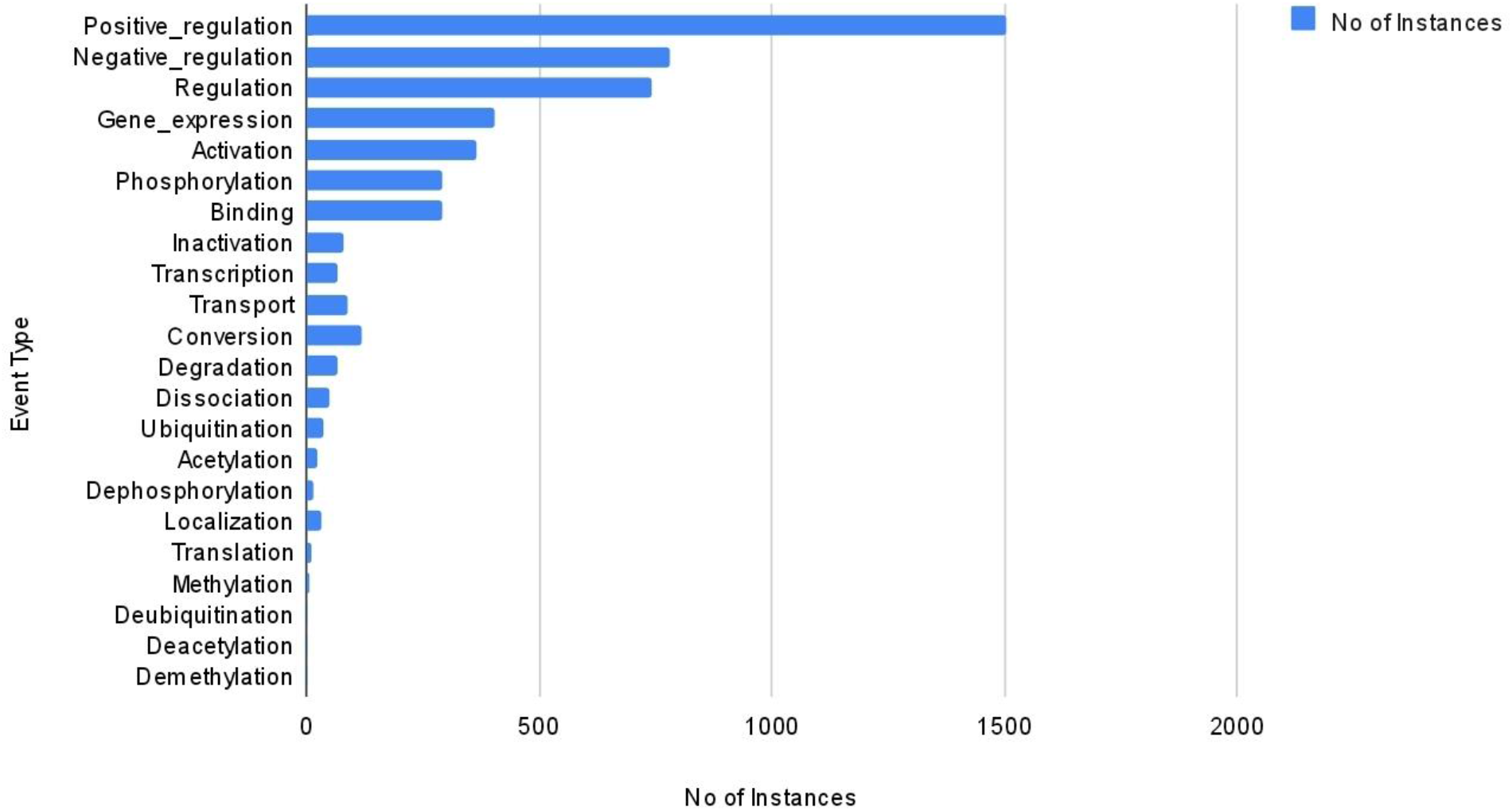
Distribution of data across the different event types

**Fig. 2.**
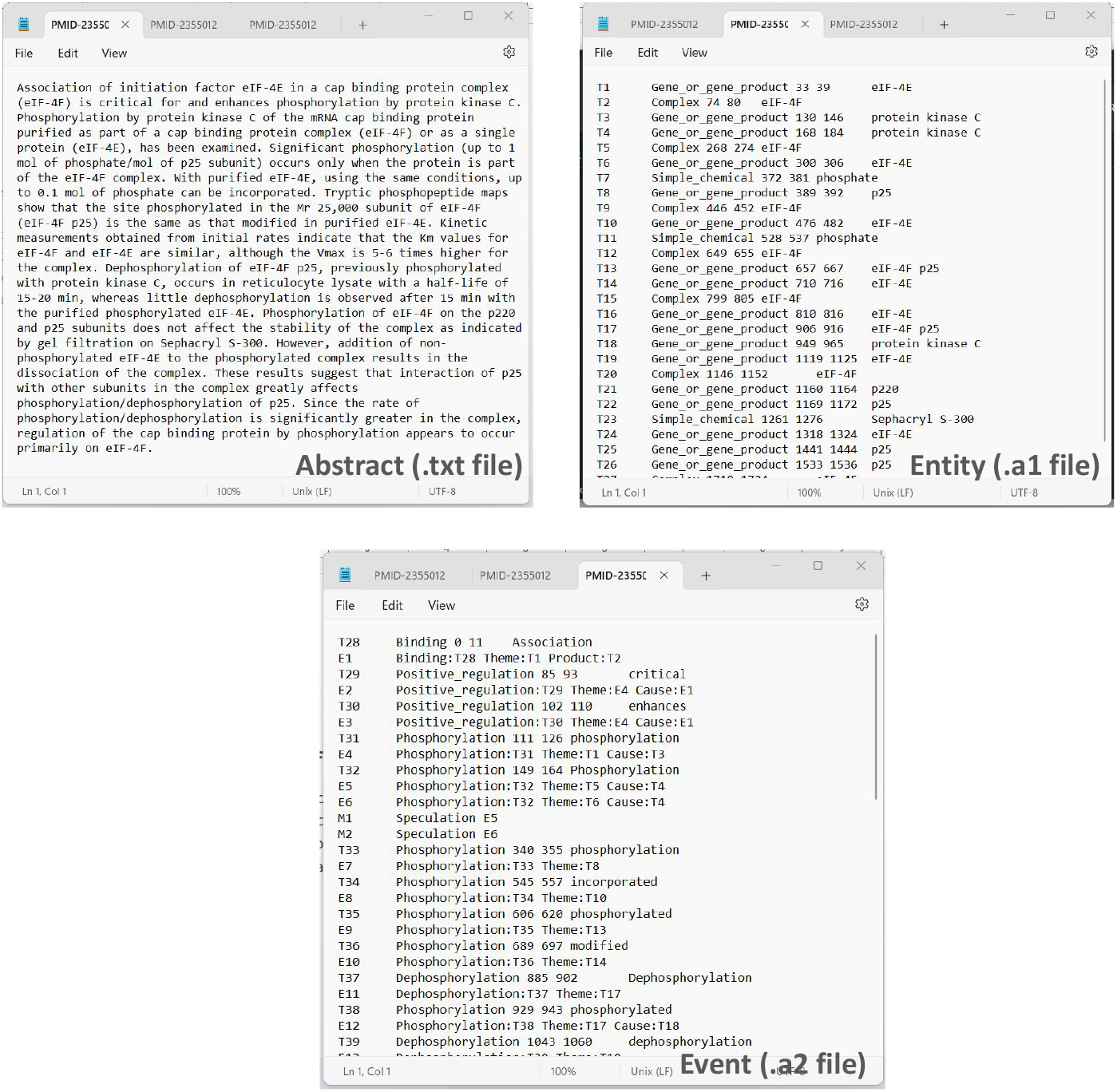
Various components of BioNLP-ST 2013 Pathway Curation Data

#### B. Data Statistics

The dataset we considered in this study comprises a total of 260 abstracts that was curated to ensure a representative cross-section of the domain. Within these abstracts, we observed an average of 19.1 reactions per abstract, cumulatively amounting to a total of 4,981 distinct reactions. Another key highlight of this dataset is the presence of 7,855 unique entities (as defined above). The event type data frequency distribution across all abstract is shown in

Fig. 1. ‘Positive Regulation’ is the most frequent event type, followed by ‘Regulation’, ‘Activation’, down to ‘Deacetylation’ which had the least representation in the dataset.

### Methodology

Our approach to tacking the problem comprises of a two-step process, systematically designed to first identify the event type of a reaction in a given statement, and then extract relevant event entities along with their roles such as Theme, Cause, Participant, etc., using the previously determined event type as shown in Fig. 3

**Fig. 3.**
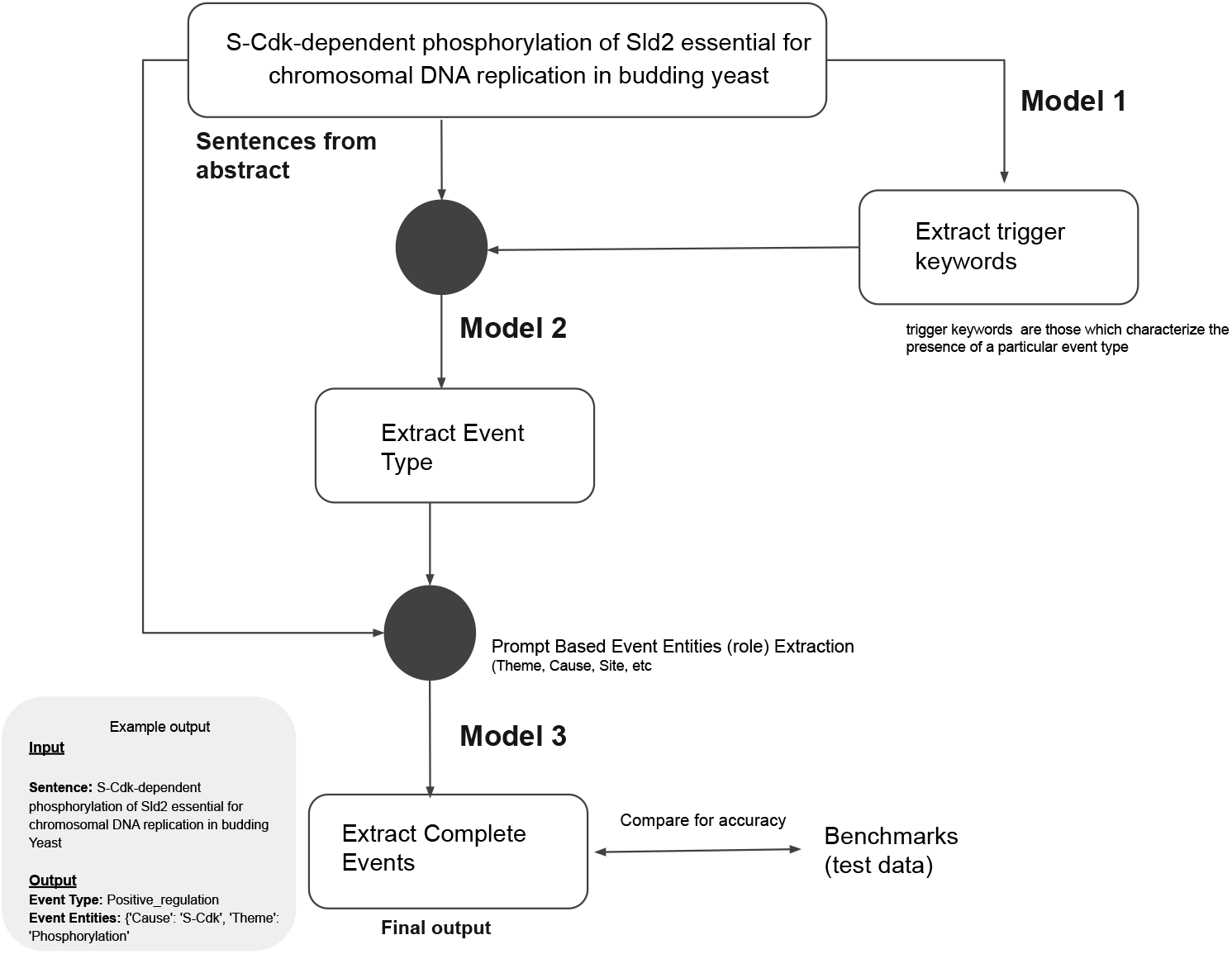
Solution strategy for tacking the Pathway Curation Task. Three separate models were first fine tuned based on GPT 3.5 and then put together as a complete pipeline.

#### Step 1: Identifying Event Type

The first step is the identification of event types from sentences describing biochemical reactions. This stage is split into two focused tasks to ensure accuracy. Initially, we train a model, which we refer to as Model 1, to detect key trigger words within a sentence. This model is fine tuned on the GPT-3.5 model with the preprocessed BioNLP-ST 2013 dataset to adapt to the task-specific format.

Subsequently, Model 2, also based on GPT-3.5, uses the triggers identified by Model 1 to determine the event type. It considers both the trigger words and the full sentence to contextualize the event. This two-tiered approach allows for a more refined extraction of event types, ensuring that the model’s output aligns closely with the complex nature of biochemical text data.

Both Model 1 and Model 2 are crucial for this identification step, having been fine-tuned from the GPT-3.5 architecture to specifically perform these tasks. Their combined use helps us achieve a high level of precision in understanding and categorizing the events. The accuracy of Model 1 and Model 2 was 91%.

#### Step 2: Extracting Event Entities with Roles

Once the event type has been identified, the subsequent step is to extract key entities associated the different roles corresponding to the event type from the abstract. These entities with roles such as Theme, Cause, Site, Participant, among others, are essential for delineating the specific dynamics of the reaction. The model, provided with both the sentence and the identified event type, employs a slot-filling method to accurately pinpoint these relevant entities.

In this phase, we first conducted experiments using a zeroshot setting to gauge the inherent capabilities of GPT-4 and GPT-3.5 without any bespoke fine-tuning. The initial results were promising (see supplementary information for the zero shot prompt), yet fine-tuning the model yielded further enhancements, this model is called Model 3 (see Fig. 3). For this fine-tuning process, we utilized GPT-3.5 as the foundational model. Employing a fill-in-the-blank strategy, the model was trained to predict specific roles (and event entities) based on the given Event Type, as outlined in Table 1. This critical step is integral to the whole process thereby significantly contributing to the efficacy of the pathway curation task. An example of the format used for fine-tuning the model is provided below:

**Table 1.** Event Types and Output Entities.

| Event Type | Reaction Types |
| --- | --- |
| Positive_Regulation | Theme & Cause |
| Negative_Regulation | Theme & Cause |
| Binding | Themes |
| Regulation | Theme, Cause & Site |
| Transcription | Theme |
| Degradation | Theme |
| Ubiquitination | Theme & Cause |
| Phosphorylation | Theme & Site |
| Localization | Theme, ToLoc, AtLoc |
| Dissociation | Theme & Product |
| Activation | Theme & Cause |
| Inactivation | Theme |
| Gene_expression | Theme |
| Transport | Theme, ToLoc, FromLoc |
| Conversion | Theme & Product |
| Dephosphorylation | Theme, Cause & Site |
| Acetylation | Theme, Cause & Site |
| Translation | Theme |

Combining Models 1, 2, and 3, we establish a comprehensive pipeline for event extraction in our pathway curation task. The process begins with Model 1 identifying trigger words in sentences. Model 2 then classifies the event type based on these triggers. Finally, Model 3 extracts detailed entities and roles associated with the identified event type. See Fig. 3 for the end to end flow. The input to this pipeline is a set of sentences, and the output is a structured set of events. This output is directly comparable with established benchmarks, allowing for an objective evaluation of the pipeline’s performance.

###### Fine-tuning Data Format

###### INPUT

For the given reaction statement, please identify the Theme & Cause for Negative_Regulation, Theme & Cause for Negative_Regulation, Themes for Binding Events in json format. Keep the json format consistent throughout the chat and do not add extra spaces or explanation outside json object.

###### Reaction Statement

Regulation of NF-kappaB RelA phosphorylation and transcriptional activity by p21(ras) and protein kinase Czeta in primary endothelial cells

###### Event Type

Negative_Regulation, Negative_Regulation, Binding

###### Entities

“Interleukin-10”:

“Gene_or_gene_product”, “inhibitor of kappaB kinase”: “Complex”, “nuclear factor kappaB”: “Complex”

###### OUTPUT

~~~
{
    “Negative_Regulation”: [
      {
           “Cause”: “Pathway”,
           “Theme”: “inhibitor of kappaB kinase”
      },
      {
           “Cause”: “Pathway”, “Theme”: “Binding”
     }],
  “Binding”: [{
           “Theme”: “nuclear factor kappaB” }]
}
~~~

### Experimental Setup

For our study, we utilized OpenAI’s API, specifically focusing on the GPT-3.5-turbo model as the base model for our model development. The first model, designed for extracting trigger keywords was fine-tuned on GPT-3.5-turbo for a total of 10 epochs. The parameters set for inference with this model were: temperature was set to 0.01 to minimize randomness in the responses, *top*_*p* was set to 1 to allow the model to consider the entire probability distribution of words, *frequency*_*penalty* was set to 0 to avoid penalizing the repetition of words, and n was set to 1 to generate a single response for each prompt. The responses from this model were formatted in JSON for ease of integration and analysis.

For the second model, which focuses on detecting the event type from the identified trigger keywords, we kept the model parameters to be the same for the sake of consistency and comparability. Similarly, the third model, which is responsible for extracting event entities with their roles, was fine tuned using the same parameters. This uniformity across all three models aids in maintaining a consistent performance metric and simplifies the process of tweaking and optimizing the models. The fine-tuning process for each of these models was completed within a time frame of 2-3 hours. All the three models were trained and tested using the PC task Data provided during the BioNLP-ST 2013 conference. For all out experiments, the train, test and validation data split were in the ratio of of 80 : 10 : 10.

## Results

Our first objective was to assess the off the shelf capabilities of GPT-3.5 and GPT-4 in understanding and classifying different event types through a zero-shot inference approach. The experiments indicated that while the models have a foundational level of comprehension for event identification, the zero-shot performance did not fully meet the accuracy standards. One notable limitation observed was the model tended to generate outputs that deviated from the set format, along with instances of partial detection of entities. The GPT-3.5 model achieved a precision of 25%, a recall of 20%, and an F1 score of 22.24%. The GPT-4 model yielded a slightly higher precision of 33.33% but had the same recall rate, and an F1 score of 25%, as detailed in Table 2.

**Table 2.**
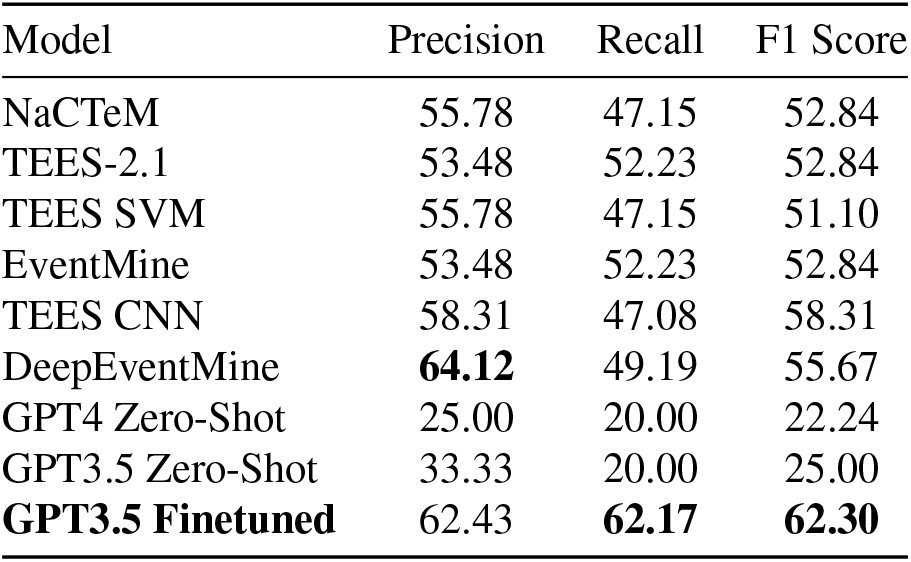
Overall accuracy of different systems for event extraction. The results of the systems were obtained from their respective papers. GPT4 Zero-Shot, GPT3.5 Zero-Shot and GPT3.5 Finetuned were the models based on LLMs

| Model | Precision | Recall | F1 Score |
| --- | --- | --- | --- |
| NaCTeM | 55.78 | 47.15 | 52.84 |
| TEES-2.1 | 53.48 | 52.23 | 52.84 |
| TEES SVM | 55.78 | 47.15 | 51.10 |
| EventMine | 53.48 | 52.23 | 52.84 |
| TEES CNN | 58.31 | 47.08 | 58.31 |
| DeepEventMine | **64.12** | 49.19 | 55.67 |
| GPT4 Zero-Shot | 25.00 | 20.00 | 22.24 |
| GPT3.5 Zero-Shot | 33.33 | 20.00 | 25.00 |
| **GPT3.5 Finetuned** | **62.43** | **62.17** | **62.30** |

Subsequent fine-tuning of the GPT-3.5 model, however, marked a substantial enhancement in performance metrics. The fine-tuned GPT-3.5 model had a precision of 62.43%, recall of 62.17%, and an F1 score of 62.30%, as shown in Table 2. The model outperformed other state-of-the-art models like TEES CNN and DeepEventMine as shown in Table 2. One thing to note was that the precision of DeepEventMine was marginally better at 64.12%.

Table 3 provides a granular view of the performance of the best performing model, fine-tuned GPT-3.5, across various event types. The model performed well for a range of event types where there were well defined contextual triggers and the reaction setting was simpler. Complex event types, such as Regulation, Positive Regulation, Negative Regulation, Conversion, etc., exhibited lower accuracy compared to other event types. These complex events, which often involve subtle linguistic cues and extensive contextual interpretation, remain challenging for automated systems. Certain event types are not included in Table 3, and this exclusion is attributed to a lack of sufficient data points of these particular events in the PC Task dataset.

**Table 3.** Performance of the best performing GPT3.5 fine-tuned model by event type. Event types not mentioned in the table did not have enough examples for training.

| Event | GPT3.5 Finetuned |  |  |
| --- | --- | --- | --- |
|  | precision | recall | F-score |
| CONVERSION | 0.5740 | 0.6326 | 0.6019 |
| PHOSPHORYLATION | 0.7818 | 0.7963 | 0.7889 |
| DEPHOSPHORYLATION | 0.6666 | 0.6666 | 0.6666 |
| ACETYLATION | 0.8571 | 0.75 | 0.7999 |
| UBIQUITINATION | 0.8235 | 0.8235 | 0.8235 |
| LOCALIZATION | 0.9166 | 0.9166 | 0.9166 |
| TRANSPORT | 0.5945 | 0.5789 | 0.5866 |
| GENE EXPRESSION | 0.7441 | 0.7272 | 0.7356 |
| TRANSCRIPTION | 1.0 | 1.0 | 1.0 |
| TRANSLATION | 1.0 | 1.0 | 1.0 |
| DEGRADATION | 0.9285 | 0.9285 | 0.9285 |
| ACTIVATION | 0.5576 | 0.5576 | 0.5576 |
| INACTIVATION | 0.875 | 0.875 | 0.875 |
| BINDING | 0.5 | 0.5 | 0.5 |
| DISSOCIATION | 0.5 | 0.5 | 0.5 |
| REGULATION | 0.5454 | 0.5393 | 0.5423 |
| POSITIVE REGULATION | 0.5585 | 0.5468 | 0.5526 |
| NEGATIVE REGULATION | 0.5384 | 0.5490 | 0.5436 |

In conclusion, the fine-tuned model based on GPT-3.5 model emerges as a clear winner for event extraction. However, the nuanced nature of more complex event types points to better model refinement strategies and dataset expansion to achieve higher coverage and eventually higher accuracy in event extraction tasks.

## Conclusions and Discussions

In conclusion, we have shown that LLMs, particularly when fine-tuned, can achieve state-of-the-art accuracy in the task of pathway curation. Although our work is an initial exploration, it lays the foundation for a more advanced application of LLMs in this domain. It also opens several areas for improvements such as using more effective prompting techniques, expanding the event specific data-sets for fine-tuning, and also fine tuning on advanced domain specific foundation models. Our results are in line with similar research efforts, such as those in (29) in their comparative analyses.

Looking forward, our works establishes that in future LLMs could play a important role for the pathway curation task. However, to realize this, the models have to be improved to better handle the intricacies of language usage in biology and the mastery of the contextual nuances for all event types.

## Supplementary Note 1: Author contribution

SKP Conceived the study. SKP and NK designed the solution strategy. NK, NK and MK conducted the experiments and generated the results. All authors wrote and reviewed the paper.

## ACKNOWLEDGEMENTS

The authors thank the ONRG Grant for the Nobel Turing challenge to The Systems Biology Institute (Grant number: N62909-21-1-2032). Nikshita Karkera was an undergraduate student intern from Thadomal Shahani Engineering College, India for 6 months at the Systems Biology Institute during the period of this research. Mahanash Kumar was an undergraduate student intern from PES University, India for 6 months at the Systems Biology Institute during the period of this research.

## Supplementary Note 2: Zero shot prompt for Model 3

#### Zero Shot Prompt SYSTEM MESSAGE

Pathway curation, an important process in molecular biology, necessitates the meticulous identification and documentation of cellular pathways, integral to understanding diverse biological functions and physiological responses. This process faces significant challenges due to the exponential growth of biomedical literature, which has outpaced traditional manual curation methods. The emergence of Generative AI offers a promising solution to these challenges, enabling efficient and accurate extraction of pathway-related information from vast quantities of scientific texts. Our study focuses on leveraging Generative AI techniques to revolutionize pathway curation, aiming to automate and enhance the extraction of intricate molecular events and interactions from biomedical research publications.

Based on the above context, please help with event extraction for the statements that will be given in subsequent messages

Before asking any questions, I want to explain you the meaning of few keywords

Theme: entity/event that undergoes the effects of the event. For example, the entity that is transcribed in a TRANSCRIPTION event or transported in a TRANSPORT event.

Cause: entity/event that is causally active in the event. Marks, for example, “P1” in “P1 inhibits P2 expression.

AtLoc, FromLoc, ToLoc : location in which the Theme entity of a LOCALIZATION event is localized (At) in LOCALIZATION events not involving movement or is transported (or moves) from/to (From/To) in LOCALIZATION and TRANSPORT events involving movement.

Site: site on the Theme entity that is modified in the event. Can be specified for modification events such as PHOSPHORYLATION.

Participant: general role type identifying an entity that participates in some underspecified way in a high-level process. Only applied for the PATHWAY type.

#### Event Type

Negative_Regulation, Negative_Regulation, Binding

#### Entities

“Interleukin-10”: “Gene_or_gene_product”, “inhibitor of kappaB kinase”: “Complex”, “nuclear factor kappaB”: “Complex”

